# Tuning in together: LSD enhances inter-brain synchrony and felt connectedness in romantic couples

**DOI:** 10.64898/2026.08.13.744639

**Authors:** NL Mason, A Czeszumski, I Totomanova, F. Trbusek, M Cavarra, S. M. Ashton, SW. Toennes, EL Theunissen, J.T. Reckweg, P. L. Lockwood, M. DeWitte, K.H. Preller, KPC Kuypers, G Dumas, P Mallaroni, JG Ramaekers

## Abstract

Social connection is fundamental to human wellbeing. Serotonergic psychedelics such as lysergic acid diethylamide (LSD) acutely heighten subjective connectedness, yet their effects on real-time social connection remain poorly understood. Using EEG hyperscanning in a randomized, double-blind, placebo-controlled crossover study, we recorded neural activity simultaneously from both members of healthy romantic couples (N=25) who received LSD (50 μg) or placebo together, across resting and interactive states. LSD increased subjective connectedness, including feelings of love, closeness, trust, and being “in sync,” while reducing loneliness, compared to placebo. This affiliative shift dissociated from the drug’s pharmacokinetic time-course, remaining elevated as subjective intensity and plasma concentration declined. In parallel, LSD increased inter-brain synchrony during shared rest, carried specifically by theta-band amplitude-envelope coupling. Importantly, this effect survived two complementary controls. First, it exceeded coupling between unrelated individuals and second the effects depended on contemporaneous neural alignment rather than shared drug-induced dynamics. Exploratory analyses showed that romantic partners with greater resting synchrony reported greater feelings of connectedness. These findings provide the first evidence that a psychedelic enhances brain-to-brain coupling between people, linking a pharmacologically induced state of felt connection to a measurable signature shared across interacting brains.

## Introduction

Social connection is fundamental to human wellbeing. The ability to form and maintain close relationships provides emotional support, shapes how people regulate affect, and contributes to psychological and physical health(1–4). Yet social connection is not generated by individuals in isolation; it emerges through ongoing interactions in which people continuously respond and adapt to one another(5,6). Understanding the mechanisms supporting this interpersonal attunement is therefore a central aim of social neuroscience.

Serotonergic psychedelics such as psilocybin and lysergic acid diethylamide (LSD) offer a potentially informative means of investigating these mechanisms: they reliably alter social experience, increasing emotional empathy, self-disclosure, and feelings of connectedness to others (for a comprehensive review see (7)). Both clinical and healthy volunteer studies further report enduring changes in perceived connectedness and in self–other boundaries that may persist beyond the acute state(8–10), leading to the hypothesis that enhanced social connection may be a key therapeutic mechanism of psychedelics(7). However, existing evidence comes from individual participants completing non-interactive social tasks. Therefore it is largely unknown whether and how psychedelics influence social processes as they unfold between people, a focus of second-person neuroscience(6).

A common way people describe feeling deeply connected to another is as “being on the same wavelength” or “sharing a mind.” Social neuroscience offers a potential substrate for these metaphors in interpersonal neural synchrony — the coupling of brain activity across individuals — which is increasingly recognised as a signature of social connection(11–13), and is associated with coordination, cooperation, empathy, and affective touch(14–20). Hyperscanning approaches, in which signals are recorded simultaneously from multiple brains, have been central to establishing this link; electroencephalography (EEG) in particular enables high-temporal-resolution measurement of inter-brain synchrony during naturalistic interaction(21). No study to date has tested whether a psychedelic modulates interpersonal neural synchrony within a dyad sharing the drug experience.

Romantic partnerships provide a particularly salient context in which to address this question. For many adults, a romantic partner is a primary source of emotional support and the relationship in which core attachment needs are met(22). Studying established couples therefore allows psychedelic effects to be examined within an existing and personally meaningful social bond. This context may also have broader clinical relevance: relationship distress and psychopathology are strongly and bidirectionally interrelated(23), motivating calls to involve partners in individual treatment(23). Indeed this approach is now beginning to be implemented in psychedelic-assisted psychotherapy(24). Romantic couples therefore provide an ecologically meaningful context in which to test whether psychedelic-induced changes in social connection extend to established interpersonal bonds.

Here we combine neuropsychopharmacology and social neuroscience to test whether the classic psychedelic LSD enhances neural and subjective signatures of interpersonal connectedness within romantic couples. Using EEG hyperscanning, we simultaneously recorded neural activity from both members of healthy romantic dyads who received either LSD (50 μg) or placebo together. We first tested whether LSD increases inter-brain synchrony relative to placebo, across resting and interactive states, validating any effects against surrogate dyads to ensure they reflect genuine, partner-specific coupling rather than shared pharmacological, sensory or environmental input. Second, we examined whether LSD enhances subjective connectedness within the dyad. Finally, in an exploratory analysis, we tested whether individual differences in LSD-induced synchrony related to individual differences in subjective connectedness.

## Methods

### Regulatory approval

This study was conducted according to the code of ethics on human experimentation established by the declaration of Helsinki (1964), amended in Fortaleza (Brazil, October 2013), in accordance with the Medical Research Involving Human Subjects Act (WMO) and was approved by the Academic Hospital and University’s Medical Ethics committee. All participants were fully informed of all procedures, possible adverse reactions, legal rights and responsibilities, expected benefits, and their right for voluntary termination without consequences. A permit for obtaining, storing, and administering LSD was obtained from the Dutch Drug Enforcement Administration. The study was registered on Onderzoekmetmensen before data collection began (https://onderzoekmetmensen.nl/nl/trial/53689; NL-OMON53689; 13-09-2022).

### Participants

Of 385 individuals assessed for eligibility, 50 were ultimately enrolled and tested (Figure S1). Those 50 healthy adults (27 women, 21 men, 2 non-binary) aged 18–40 years (mean ±SD: 25.46 ±5.25) formed 25 romantic couples (21 mixed-gender couples, 3 female same-gender couples, one non-binary couple). Participants were recruited via word of mouth and advertisements distributed on social media. Eligible couples were required to have been in a relationship for at least 6 months (44% more than 2 years, 40% between 1 and 2 years, 16% between 6 months and 1 year). All participants had to have had at least one previous lifetime exposure to a classic psychedelic (e.g., psilocybin, mescaline, LSD), but no psychedelic use within the 3 months preceding enrolment (mean ±SD of lifetime psychedelic use: 9.52 ±12.60). Further inclusion and exclusion criteria are described in the Supplementary Materials and Table S1.

### Study design

Participants were enrolled between December 2022 and February 2026 in a double-blind, placebo-controlled, two-way crossover study conducted at Maastricht University, the Netherlands. Each romantic dyad participated in two experimental sessions, during which both partners received either an oral dose of LSD (50 μg) or placebo, with both members of the couple always receiving the same treatment on a given test day. The 50 μg dose was selected because it reliably induces a psychedelic state while still allowing participants to interact and engage in interpersonal tasks. The order of treatments (LSD–placebo or placebo–LSD) was randomized and balanced using an independent centralized randomisation procedure generated by researchers who had no contact with participants. Sessions took place in an ecologically valid, living room–style laboratory setting, with partners seated across from one another at a table to support naturalistic interaction. Sessions were separated by a minimum washout period of 14 days to avoid carry-over effects. EEG hyperscanning and measures of connectedness and drug intoxication were performed during the peak drug effects, starting 1.5 hours after drug administration and continuing until 7 hours post-dose (Figure 1).

**Figure 1.**
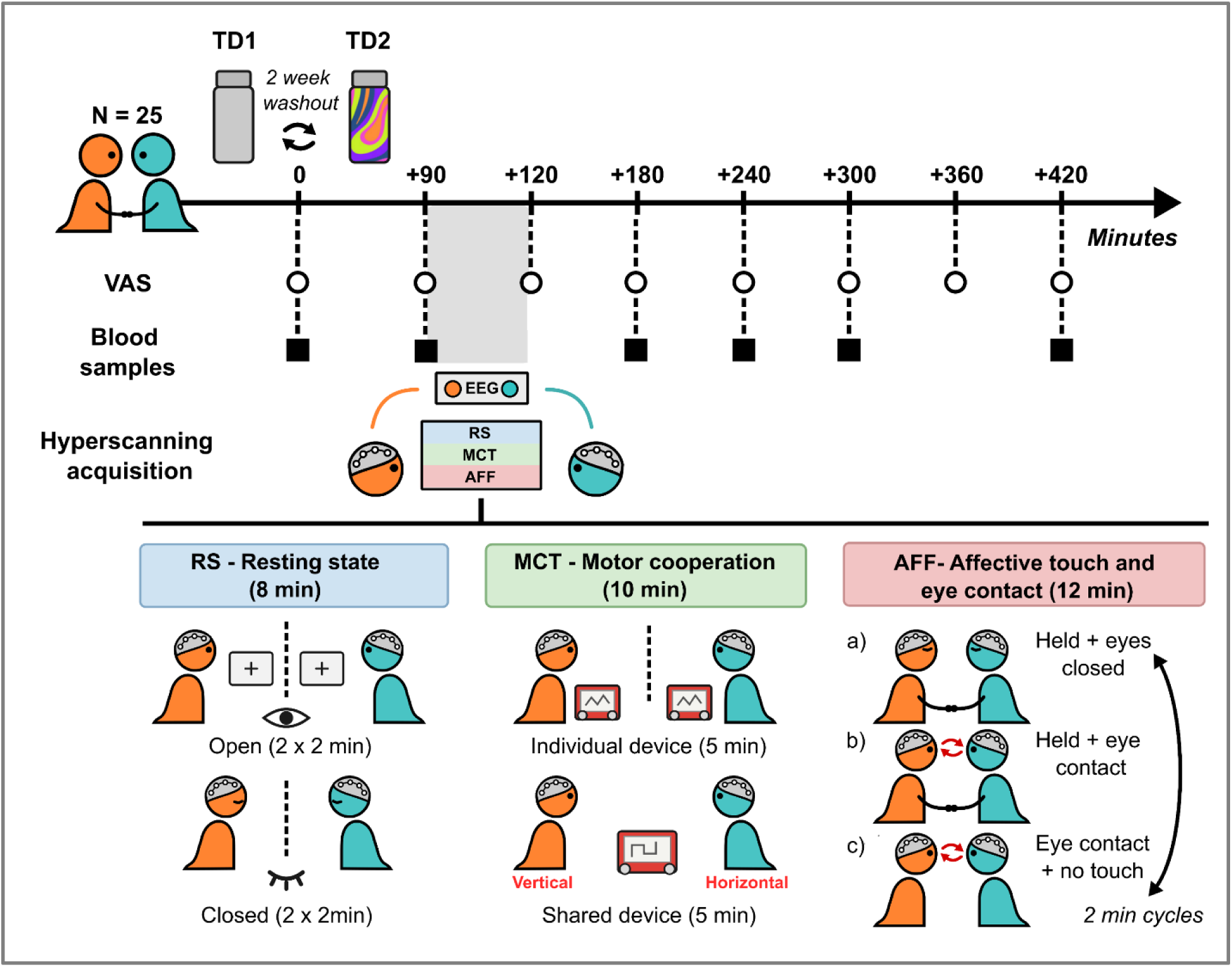
Schematic overview of the experimental timeline. Timeline of procedures on each test day, shown relative to drug administration (t = 0). After arrival, participants completed baseline blood samples, questionnaires and visual analogue scales (VAS), followed by drug administration (LSD 50 μg or placebo). EEG hyperscanning was conducted during the peak drug window and included the resting-state (RS), motor cooperation task (MCT), and affective touch and eye contact paradigm (AFF) recordings. Questionnaires and VAS assessing social feelings and drug effects were administered at predefined time points across the day. Five venous blood samples for pharmacokinetic analyses were collected after drug administration and aligned with the EEG tasks and questionnaire blocks. The procedure was identical on LSD and placebo days, with at least 14 days between sessions.

### EEG

Neural activity was recorded simultaneously from both members of the couple using 32-channel EEG (BrainAmp) caps arranged according to the international 10–20 system. A linked-mastoid reference and a forehead ground (AFz) were used for all recordings. Horizontal and vertical electro-oculogram (EOG) channels were applied to monitor eye movements and blinks. Cardiac activity was recorded using a three-electrode electrocardiogram (ECG). For most tasks, participants rested their chins on a padded chin rest to minimise head movements and associated muscle artefacts. Before electrode placement, the skin was cleaned with alcohol and lightly abraded with conductive gel to reduce impedance. EEG and EOG signals were sampled at 500 Hz. Throughout each session, neuroelectric activity in both participants was continuously recorded while they engaged in the experimental tasks and naturalistic interactions.

### Resting-state (RS)

Participants underwent an 8-minute resting-state paradigm with alternating 2-minute blocks of (a) eyes open while fixating a central cross and (b) eyes closed. Each condition was presented twice, for a total of 4 minutes per condition. During the scan, the participants were separated from each other by a visual barrier, which prevented eye contact between partners.

### Motor cooperation task (MCT)

To probe cooperative motor behaviour, couples performed a motor cooperation task adapted from prior hyperscanning work(20). In this task, participants were instructed to draw predefined shapes (e.g., an “x”, a house, a vase) using a mechanical drawing toy (Etch A Sketch®). First, they drew these shapes individually, each with their own Etch A Sketch, while separated by a visual barrier. The barrier was then removed and they were instructed to draw different predefined shapes together. In the joint condition, they shared a single mechanical drawing toy with two control knobs, one governing vertical and the other horizontal movement of a single drawing stylus. Partners were seated facing each other, and each drawing session (individual and joint) lasted 5 minutes. They were not allowed to speak during the task. On each test day, couples drew different images than on the other dosing day to reduce practice effects.

### Affective touch and eye contact (AFF)

To assess neural synchrony during non-verbal social contact, couples completed an affective touch and eye contact paradigm validated for hyperscanning(17,25). Partners sat facing each other and completed three conditions: (a) holding their dominant hands while keeping their eyes closed, (b) holding their dominant hands while maintaining eye contact, and (c) maintaining eye contact without physical touch. Each condition lasted 2 minutes and was repeated twice in a fixed sequence, yielding a total of 4 minutes per condition (12 minutes in total).

### Preprocessing

Dual-participant EEG was recorded in a single 70-channel BrainVision file (32 EEG channels per participant plus vertical EOG, horizontal EOG, and ECG; 500 Hz, INT_16 format). Files were processed offline using MNE-Python. Each dyadic recording was first split into two independent single-participant datasets by partitioning channels on the basis of their naming convention. Immediately after splitting, all recordings were resampled from 500 Hz to 250 Hz to reduce memory overhead prior to artifact rejection. Standard electrode coordinates were assigned using the MNE standard_1020 montage after correcting a capitalisation inconsistency in the original BrainVision file (PZ → Pz).

Each participant’s data were then filtered using a zero-phase finite impulse response (FIR) filter. A notch filter was applied at 50 Hz and 100 Hz to suppress European mains noise and its first harmonic, followed by a bandpass filter from 0.1 to 40 Hz (high-pass cutoff 0.1 Hz; low-pass cutoff 40 Hz).

Automated bad channel detection was performed using the RANSAC algorithm as implemented in the PyPREP library (random seed 1337; sample proportion 0.25; correlation threshold 0.75; bad fraction threshold 0.40; channel-wise mode). RANSAC was applied to EEG channels only, excluding EOG and ECG. Detected bad channels were marked but not yet removed at this stage. Continuous data were then subjected to Artifact Subspace Reconstruction (ASR; cutoff = 5.0 SD) to suppress high-amplitude transient artifacts while preserving slower brain dynamics.

Independent component analysis (FastICA algorithm; random seed 42) was applied to the ASR-cleaned data. Components were automatically identified as artifactual based on their correlation with the EOG channels (ocular artifacts; up to three components rejected per participant) or the ECG channel (cardiac artifacts; up to two components rejected per participant). Identified components were subtracted from the data. RANSAC-flagged bad channels were then reconstructed by spherical spline interpolation. Finally, all EEG channels were re-referenced to the average of all scalp electrodes.

### Channel Harmonisation

A critical methodological concern was that LSD administration produced significantly more RANSAC bad channels than placebo across participants. Without correction, this would introduce a systematic confound: inter-brain synchrony (IBS) computed from different channel subsets across conditions would be incomparable within the same dyad. To address this, a per-dyad channel harmonisation procedure was applied prior to IBS computation. For each dyad, the union of all RANSAC-flagged bad channels was assembled across all four data cells contributing to that dyad’s contrast (both participants under both drug conditions). This union set was then excluded from IBS computation in every cell for that dyad, ensuring that all within-dyad comparisons were computed on a strictly identical channel set. In addition, the channel A2 (right mastoid, retained as an EEG channel during preprocessing but not a cortical source) was excluded universally from all IBS computation.

### Behavioral and subjective experience

#### Visual Analogue Scale

Visual analogue scales (VAS) were administered at predefined time points (Figure 1) throughout each testing day to assess subjective feelings of social behaviour and drug effects. VAS items were presented as 100-mm horizontal lines anchored with “not at all” on the left and “extremely” on the right. For social VAS items, the midpoint (50 mm) was explicitly labelled as representing the participant’s “normal” level for that construct. Participants indicated their responses by placing a mark on the line, and scores were quantified as the distance in millimetres from the left anchor. Social VAS items included: sociable, loving, lonely, connected, safe, trusting, commitment, attachment, close, in sync, attractiveness. Drug effect items included: under drug influence, good drug effect, bad drug effect, I like the drug, feeling high. In addition to being analysed separately, relational and drug-effect items were combined into Connectedness and Drug-intensity composites, respectively (see Statistical analysis).

#### Plasma LSD pharmacokinetics

Six blood samples (1×10 mL, Lithium Heparine) to determine LSD plasma concentration were collected throughout the testing day (Figure 1). See Supplement for information on analysis procedures and results.

### Statistical analysis

#### Subjective outcomes and brain-behavior associations

Statistical analysis of behavioral and subjective outcome measures was conducted in IBM SPSS Statistics 28 using a Linear Mixed Model (LMM) analysis. For the VAS measures, separate LMMs were run with Treatment (LSD or placebo), Time (8 assessment points) and their interaction (Treatment*Time) as fixed effects. Because both members of each couple were dosed together and completed each session jointly, observations were treated as nested within couples: random intercepts were included for couple and for participant within couple. A first-order autoregressive (AR1) covariance structure was specified for the repeated measurements across time within each treatment session. Models were estimated using restricted maximum likelihood (REML). For the drug-experience items the participant-within-couple variance was estimated at zero, so this level was omitted; the couple-level intercept was retained. To account for multiple comparisons across the VAS items, *p*-values were corrected using the Benjamini–Hochberg false discovery rate procedure, applied separately within the relational item family and the drug-experience item family.

In order to assess whether LSD-induced changes in neural synchrony were related to subjective feelings measured via the VAS, we constructed composite scores on both the neural and self-report sides of the analysis, limiting the number of statistical comparisons and deriving theoretically meaningful measures. Because each VAS item was rated individually by both partners but neural synchrony is a dyadic measure, partner ratings were averaged to yield a single couple-level score per item.

For the relational self-report items, we conducted an exploratory principal component analysis (PCA) on the couple-level relational VAS items. Sampling adequacy was acceptable (Kaiser-Meyer-Olkin = .84; Bartlett’s test of sphericity p < .001). The analysis yielded a dominant first component (56% of variance) on which eight items loaded strongly (loadings .67–.95; Table S2): loving, connected, commitment, attachment, close, in-sync, attractive, and trusting. These were averaged into a Connectedness composite which showed high internal consistency (Cronbach’s α = .94). Subjective drug intensity was indexed by a separate composite of four drug-experience items. A PCA of the five drug items yielded a dominant first component (83% of variance) on which four items loaded strongly (drug influence, good drug effect, drug liking, and feeling high; loadings .95–.98; Table S3); these were averaged into a Drug-intensity composite (Cronbach’s α = .96). As all VAS items shared a common 0–100 scale, composites were computed as unstandardized item means.

For the neural analyses, we considered the connectivity metrics and region of interest (ROI) pairs that showed a significant treatment-related effects; all four amplitude envelope correlation (AEC) ROI pairs and both weighted phase-lag index (wPLI) ROI pairs met this criterion. For each ROI pair, eyes-open and eyes-closed resting-state values were first averaged to yield a single resting-state estimate per dyad and treatment condition. Within each metric the ROI-pair measures were highly intercorrelated, so they were standardized (z-scored across the full sample) and averaged into an AEC composite (α = .98) and a wPLI composite (α = .83). The two metrics were analyzed separately, as phase and amplitude coupling index distinct connectivity phenomena. For the primary brain–behavior analysis, Connectedness and Drug-intensity scores were taken from the VAS assessment immediately adjacent to the resting-state recording. Spearman rank correlations were then computed between the two inter-brain synchrony composites and the two VAS composites (Connectedness, drug intensity) within the LSD condition (N = 23). Given the modest sample size, rank-based correlations were used to reduce sensitivity to non-normality and influential observations; effect sizes (Spearman’s ρ) are reported with percentile bootstrap 95% confidence intervals (5,000 resamples), both uncorrected and with Benjamini-Hochberg false discovery rate (FDR) correction across the four tests. To assess whether associations with Connectedness were also present outside the LSD state, corresponding correlations with Connectedness were examined under placebo (Supplementary Material).

### Inter-Brain Synchrony Analysis and Statistics

#### Connectivity Metrics and Frequency Bands

Inter-brain synchrony was quantified using two complementary connectivity metrics: the weighted phase-lag index (wPLI) and amplitude envelope correlation (AEC). wPLI indexes phase synchronisation between participants while discounting zero-lag co-fluctuations that may reflect volume conduction or common reference artifacts. For inter-brain connectivity, it removes spurious synchronization associated with common environmental input like noise, thus focusing on synchronization that is more related to the ongoing exchange of information(26). AEC quantifies co-modulation of broadband amplitude envelopes after bandpass filtering, capturing slower envelope-level coupling that is theoretically independent of phase relationships(27,28). Both metrics were computed on non-overlapping 2-second epochs (minimum 60 non-overlapping 2-second epochs, i.e. 120 s, of retained data per cell required). For wPLI, the cross-spectral density was estimated epoch-by-epoch and accumulated to yield a single wPLI estimate per recording. For AEC, the signal was bandpass filtered using a 4th-order Butterworth filter, the amplitude envelope was extracted via the Hilbert transform, and Pearson correlation was computed between participant envelopes epoch-by-epoch and averaged. Because IBS is measured between two separate individuals, orthogonalisation was not applied for AEC. Analysis were restricted to theta (4–8 Hz), alpha (8–13 Hz), and beta (13–30 Hz); delta and gamma were not examined because they are especially vulnerable to eye-movement and muscle artefacts in datasets involving naturalistic movement.

#### Region-of-Interest Aggregation

Electrode-level connectivity matrices were first mapped onto a common 31-channel template (the union of all ROI-constituent channels) to handle per-dyad channel exclusions while maintaining a consistent spatial reference frame. Values at excluded electrodes were set to NaN. Template matrices were then aggregated into five bilateral anatomical regions of interest (ROIs): Frontal (AF3, AF4, F7, F3, FZ, F4, F8), Fronto-central (FT7, FC5, FC1, FC2, FC6, FT8), Central (T7, C3, C4, T8), Parieto-temporal (TP7, CP5, CP1, CP2, CP6, TP8, P7, P3, Pz, P4, P8), and Occipital (O1, Oz, O2). For each band, ROI-pair values were computed as the mean of all electrode-pair values within the corresponding ROI block, symmetrised across the upper and lower triangle. This yielded 15 unique ROI pairs (5 × 5 upper triangle including the diagonal) per metric per band.

#### Hierarchical Statistical Design

We adopted a hierarchical two-pass testing strategy to reduce the total multiple comparison burden while retaining sensitivity to condition-specific drug effects. The analysis included three tasks: Resting State (RS), Affective Touch (AFF), and Motor Cooperation Task (MCT).

#### Omnibus tests

For each task, metric, frequency band, and ROI pair, two omnibus tests were conducted: (1) a Drug main effect (LSD vs. Placebo, averaged across conditions) and (2) a Drug × Condition interaction. For RS and MCT, the interaction was operationalised as the difference in within-participant LSD–Placebo deltas across the two conditions (Eye-State EO vs. EC for RS; Alone vs. Together for MCT). For AFF, the Drug × Touch Condition interaction (across three touch conditions: HH-EO, HH-EC, NH-EO) was tested with a permutation F-test on per-dyad LSD–Placebo deltas, replacing the pairwise t-test approach. All omnibus tests used sign-flip permutation testing (10,000 permutations) on paired differences, yielding a two-tailed empirical p-value. False discovery rate (FDR) correction was applied within each task × metric family using the Benjamini–Hochberg procedure (q < 0.05; 45 tests per contrast per family (3 frequency bands × 15 ROI pairs, pooled)). In total, 540 omnibus tests were conducted (3 tasks × 2 contrasts × 2 metrics × 3 bands × 15 ROI pairs). Gating simple-effect tests behind a significant interaction avoided unconditionally testing all 630 possible simple-effect comparisons (3 tasks × up to 3 conditions per task × 2 metrics × 3 bands × 15 ROI pairs), restricting follow-up testing to interaction-driven effects only.

For the Drug main-effect contrast, each dyad’s inter-brain synchrony was first normalised against a cross-couple surrogate distribution within the same drug condition prior to testing. This normalisation controls for non-specific, drug-induced increases in overall signal similarity — for example, LSD homogenising spectral content — which would inflate inter-brain synchrony even between unrelated individuals. The Drug × Condition interaction contrasts were not surrogate-normalised in this way, as they test condition-specific differences in the drug effect rather than its overall magnitude

#### Gated simple effects

For each ROI pair where the omnibus Drug × Condition interaction survived FDR correction, follow-up simple-effect contrasts were run comparing LSD vs. Placebo within each individual condition. Simple effects were likewise tested with sign-flip permutation t-tests (10,000 permutations) and FDR-corrected within the gated set per task × metric. As with the Drug main-effect contrast, each dyad’s inter-brain synchrony within the gated simple-effect tests was first z-scored against a cross-couple surrogate null specific to that drug condition and individual task condition (e.g., LSD–Eyes-Closed against its own surrogate distribution), prior to testing. This protected approach ensures that simple effects are only interpreted in the context of a significant interaction.

Effect sizes are reported as Cohen’s *d* computed from within-pair differences, with 95% confidence intervals estimated via percentile bootstrap resampling (1,000 resamples).

#### Surrogate Pair Validation

To establish whether observed IBS values reflected genuine inter-brain coupling above chance levels, a surrogate analysis was conducted for all ROI pairs that survived FDR correction in either the omnibus or gated simple-effect tests(29). For each significant (task, metric, band, ROI pair), 1000 independent random cross-couple pairings were generated per drug × condition cell by reassigning participant EEG recordings from different dyads. IBS was recomputed for each surrogate pairing and averaged to obtain a surrogate null distribution mean and standard deviation. The real group mean IBS for each drug was compared against its surrogate null distribution using an empirical, one-tailed p-value (Phipson & Smyth, 2010): p = (n_exceed + 1) / (N_surrogates + 1), where n_exceed is the number of surrogate permutation means at or above the real mean. A corresponding z-score [(real mean − surrogate mean) / surrogate SD] was also computed for effect-size reporting. An above-surrogate flag was assigned where p < 0.05.

## Results

### LSD produces robust, time-varying subjective drug effects

LSD produced marked subjective drug effects that varied across the testing day. Ratings (VAS) of Being Under the Influence (*F*_(7, 512.87)_ = 74.33, *p* < 0.001), Good Drug Effect (*F*_(7, 512.99)_ = 63.93, *p* < 0.001), Bad Drug Effect (*F*_(7, 510.98)_ = 6.82, *p* < 0.001), Drug Liking (*F*_(7, 542.44_) = 51.36, *p* < 0.001) and Feeling High (*F*_(7, 514.27)_ = 81.36, *p* < 0.001) each showed a Treatment × Time interaction. Ratings were higher under LSD compared to placebo, with the magnitude of these differences varying across timepoints (Figure 2; Figure S2). All items survived FDR correction (all q ≤ .001), see Table S4 for effect sizes.

**Figure 2.**
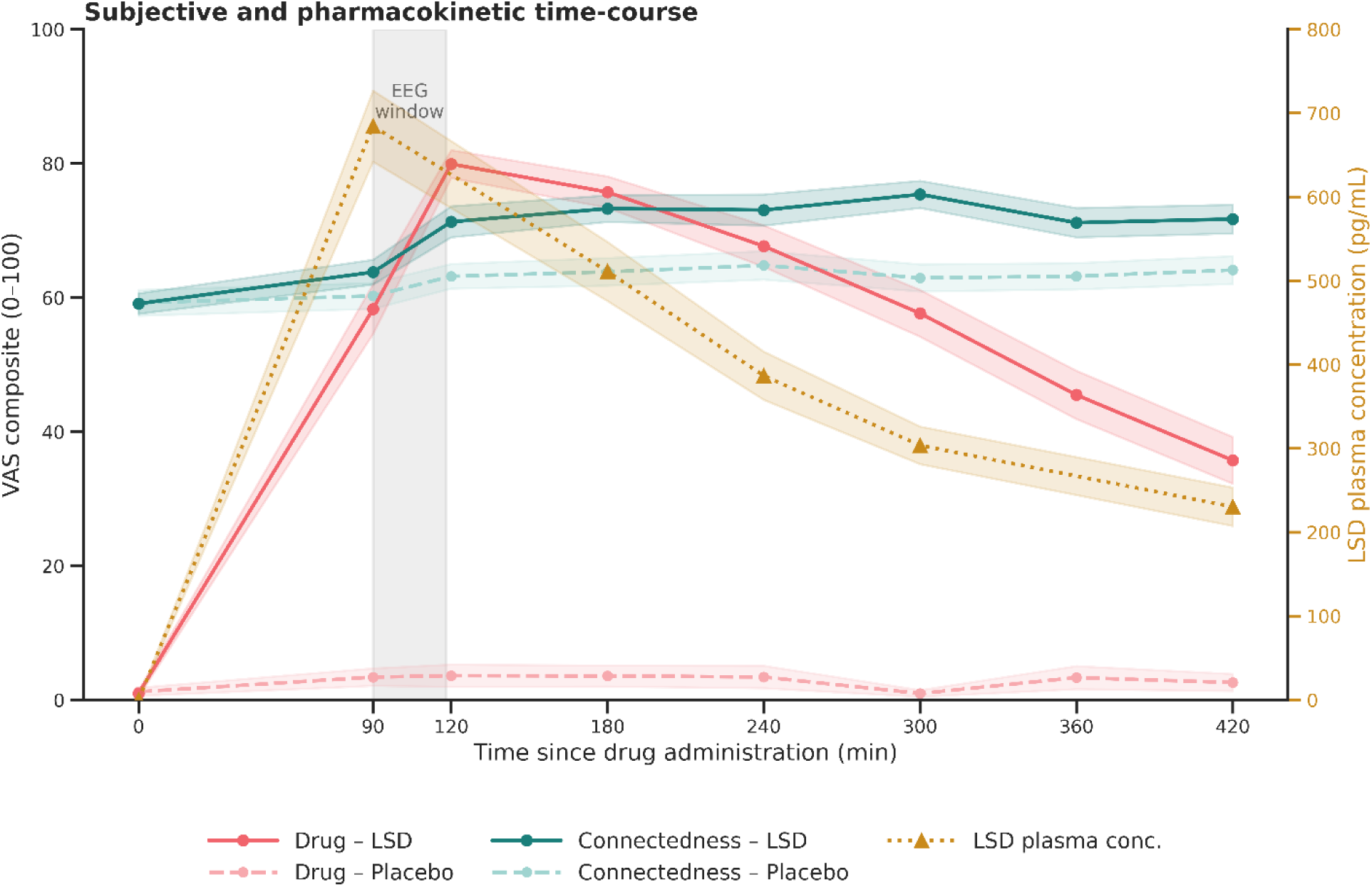
Time-course of subjective effects and LSD plasma concentration across the experimental session. Mean trajectories of the Connectedness composite (teal) and Drug-intensity composite (coral) are shown for the LSD (solid) and placebo (dashed) conditions, based on visual analogue scale (VAS) ratings from individual participants at eight timepoints from 0 to 420 min relative to drug administration. The Connectedness composite comprises eight relational items and the Drug-intensity composite four drug-experience items (see Methods). Mean LSD plasma concentration (gold, dotted; right axis) is overlaid for reference (See supplementary results). Shaded bands denote ±1 standard error of the mean. The grey region marks the EEG recording window (∼90–118 min post-dose). Under LSD, subjective drug intensity closely tracked plasma concentration, both peaking early and declining thereafter, whereas Connectedness rose and remained elevated across the session; both measures stayed near baseline under placebo.

### LSD increases affiliative and perceived interpersonal connection

LSD broadly increased affiliative feelings, including Connectedness, Attachment, Commitment, Safety, Closeness, Attractiveness, Loving, Trusting, and feeling In Sync, while reducing Loneliness. Connectedness, Safety, Commitment, Attachment, Closeness, and Attractiveness each showed Treatment × Time interactions (Connected: *F*_(7, 433.76)_ = 3.23, *p* = .002; Safe: *F*_(7, 463.24)_ = 2.53, *p* = .015; Commitment: *F*_(7, 480.38)_ = 3.40, *p* = .001; Attachment: *F*_(7, 479.69)_ = 3.85, *p* < .001; Close: *F*_(7, 454.00)_ = 2.31, *p* = .025; Attractiveness: *F*_(7, 474.56)_ = 2.39, *p* = .021), reflecting higher ratings under LSD than placebo, with the magnitude of these differences varying across the session. Ratings of Loving, Trusting, and In Sync were also higher under LSD, with these differences remaining consistent across time (main effects of Treatment: *F*_(1, 113.56)_ = 31.68, *F*_(1, 122.83)_ = 32.20, and *F*_(1, 144.33)_ = 37.95, respectively; all *p* < .001; Treatment × Time: all *F* ≤ 1.74, all *p* ≥ .098). Loneliness was lower under LSD than placebo (*F*_(1, 117.71)_ = 20.58, *p* < .001), without variation across time (*F*_(7, 453.14_) = 0.56, *p* = .79). Sociability did not differ between treatments (Treatment: *F*_(1, 125.39)_ = 0.94, *p* = .34; Treatment × Time: *F*_(7, 440.65)_ = 1.43, *p* = .19). All significant effects survived FDR correction (all *q* ≤ .028; Fig. 2 and Fig. S3), with effect sizes reported in Table S4.

To summarize the temporal dynamics, the relational ratings were combined into a Connectedness composite and the drug-experience ratings into a Drug-intensity composite. Under LSD, Drug-intensity rose and fell in close correspondence with plasma LSD concentration (See supplementary results), whereas Connectedness increased and remained elevated across the session (Fig. 2). Thus, the increase in interpersonal connectedness persisted even as subjective drug intensity and plasma LSD concentrations declined.

### LSD selectively enhances surrogate-validated inter-brain coupling during rest

Twenty-two dyads contributed to the resting-state (RS) analysis, 23 to affective touch (AFF), and 18 to motor cooperation (MCT), after excluding dyads with missing sessions or insufficient artifact-free data (<120 s per cell). Effects are reported at BH-FDR–corrected significance within each task × metric family (q < 0.05).

Across tasks, LSD altered inter-brain coupling in a frequency- and state-dependent manner, with the clearest surrogate-validated effects emerging during resting state; descriptive patterns across all tasks, frequency bands, and connectivity metrics are shown in Figure S4 and S5.

The clearest partner-specific effect of LSD emerged during rest, where inter-brain theta AEC was consistently higher under LSD than placebo across four region-of-interest (ROI) pairs, with the magnitude of this difference varying by eye state (Drug × Eye State: Central–Central, *t*(21) = −3.04, *d* = −0.65; Fronto-central–Central, *t*(21) = −3.19, *d* = −0.68; Fronto-central–Parieto-temporal, *t*(21) = −3.45, *d* = −0.74; Central–Parieto-temporal, *t*(21) = −3.58, *d* = −0.76; all *q* < .05; Fig. 3). The LSD–placebo difference was larger during eyes closed than eyes open; the negative interaction statistics therefore reflect the coding of the eye-state contrast rather than a reduction in theta AEC under LSD. Gated simple-effect tests within each eye-state condition did not survive correction (all *q* > .10), indicating that the eye-state-dependent interaction, rather than either simple effect alone, characterized the theta-AEC response to LSD (Figure 3; Figure S4).

**Figure 3.**
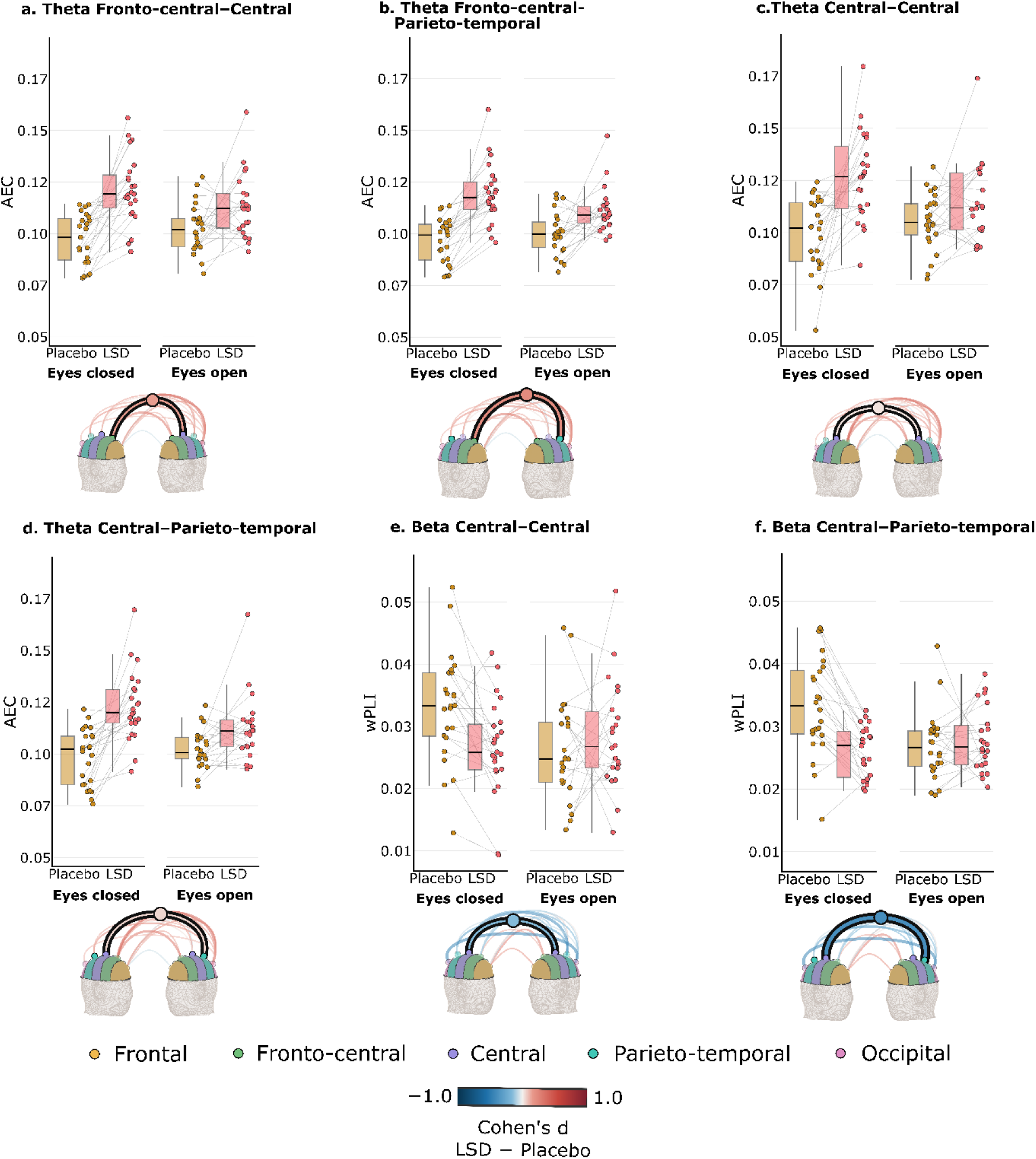
Region-specific inter-brain synchrony during resting state (RS). Inter-brain synchrony (IBS) is shown for selected region-of-interest (ROI) pairings among the electrode clusters, with each panel corresponding to the significant ROI-ROI pairings which survived the surrogate validation, within a given frequency band and connectivity metric. Panels **(a–d)** display theta-band amplitude envelope correlation (AEC) for the Central–Central, Fronto-central–Central, Fronto-central–Parieto-temporal, and Central–Parieto-temporal pairings, respectively; panels **(e–f)** display beta-band weighted phase-lag index (wPLI) for the central–central and central–parieto-temporal pairings. Within each panel, data are split by drug condition (placebo, gold; LSD, coral) and by eye condition (eyes closed, EC; eyes open, EO). Brain renders depict the ROI-ROI pairings which showed significant differences under LSD versus placebo, with the pairings which survived the surrogate validation indicated in black. Boxplots depict the median and interquartile range (IQR); whiskers extend to the furthest data points within 1.5× IQR of the quartiles; overlaid points represent individual dyads, and thin grey lines connect each dyad across the placebo and LSD sessions within a given eye condition.

To test whether significant inter-brain effects exceeded coupling expected from unrelated individuals, real dyadic synchrony was compared against a null distribution of surrogate dyads formed by pairing partners across different couples within the same session, drug, and task condition (1,000 pairings per cell; (30)). All four theta-AEC pairs exceeded the surrogate distribution under LSD (*z* = 2.86–4.02, all empirical *p* ≤ .005), whereas under placebo, theta AEC exceeded the surrogate distribution only at Central–Central (*z* = 1.85, *p* = .039). A complementary within-couple temporal-shift surrogate, which preserved the shared pharmacological context while disrupting fine-grained temporal alignment, further showed that theta-AEC coupling at all four ROI pairs depended on contemporaneous amplitude-envelope alignment under both LSD and placebo (all empirical p = .001). Together, these controls indicate that the resting-state effects reflected partner-specific, temporally aligned inter-brain coupling rather than general drug-induced similarity or shared nonstationary dynamics.

LSD also altered the eye-state dependence of beta-band phase synchrony (wPLI), rather than uniformly increasing or decreasing it. Drug × Eye State interactions emerged at Central–Central (t(21) = 3.47, *d* = 0.74, *q* = .0338) and Central–Parieto-temporal (t(21) = 4.45, *d* = 0.95, *q* = .0090; Fig. 3). Under placebo, eye closure increased beta wPLI relative to eyes open, whereas under LSD this pattern was abolished and slightly reversed. Accordingly, gated simple effects showed lower beta wPLI under LSD than placebo during eyes closed (Central–Parieto-temporal: t(21) = −3.74, *d* = −0.80, *q* = .0040; Central–Central: t(21) = −2.43, *d* = −0.52, *q* = .0464; Figure S5), with no significant differences during eyes open. Both beta-wPLI ROI pairs nevertheless exceeded their cross-couple surrogate distributions under LSD (z = 3.47–3.57, all p ≤ .002), but not under placebo.

In contrast, the structured interactive tasks showed no robust surrogate-validated effects. During affective touch, the effect of LSD on inter-brain alpha AEC varied across touch conditions at four ROI pairs (Drug × Touch Condition: Fronto-central–Fronto-central, F = 12.82, q = .0210; Frontal–Occipital, F = 10.81, q = .0210; Central–Occipital, F = 8.20, q = .0210; Fronto-central–Occipital, F = 10.81, q = .0225). Among the gated simple-effect contrasts, only Central–Occipital coupling during hand-to-hand contact with eyes open was significantly higher under LSD than placebo (t(22) = 4.59, d = 0.96, q < .001). However, no affective-touch effect exceeded the surrogate distribution under either LSD or placebo (all p > .43), and no wPLI effects survived correction. During motor cooperation, no effects survived FDR correction for the prespecified Drug × Condition interaction comparing individual and joint drawing. Thus, surrogate-validated effects were selectively observed during rest rather than during the structured interactive tasks.

### Resting inter-brain amplitude coupling, but not phase synchrony, relates to subjective connectedness under LSD

In an exploratory analysis, greater resting inter-brain amplitude coupling under LSD was associated with greater subjective Connectedness (Spearman’s ρ = .49, 95% CI [.09, .78], *p* = .018; Fig. 4). However, this association did not survive FDR correction across the four tested associations (*q* = .072) and should therefore be interpreted cautiously. Resting amplitude coupling was also negatively associated with Drug-intensity (ρ = −.44, 95% CI [−.75, .04], *p* = .037, *q* = .074), whereas phase synchrony was unrelated to either Connectedness (ρ = −.02, *p* = .939) or Drug-intensity (ρ = −.17, *p* = .433; both *q* ≥ .577). The association between amplitude coupling and Connectedness was absent under placebo (ρ = −.05, *p* = .802), suggesting that it was specific to the LSD state rather than a stable characteristic of the couples. Further exploratory analyses of its temporal specificity are reported in the Supplementary Results.

**Figure 4.**
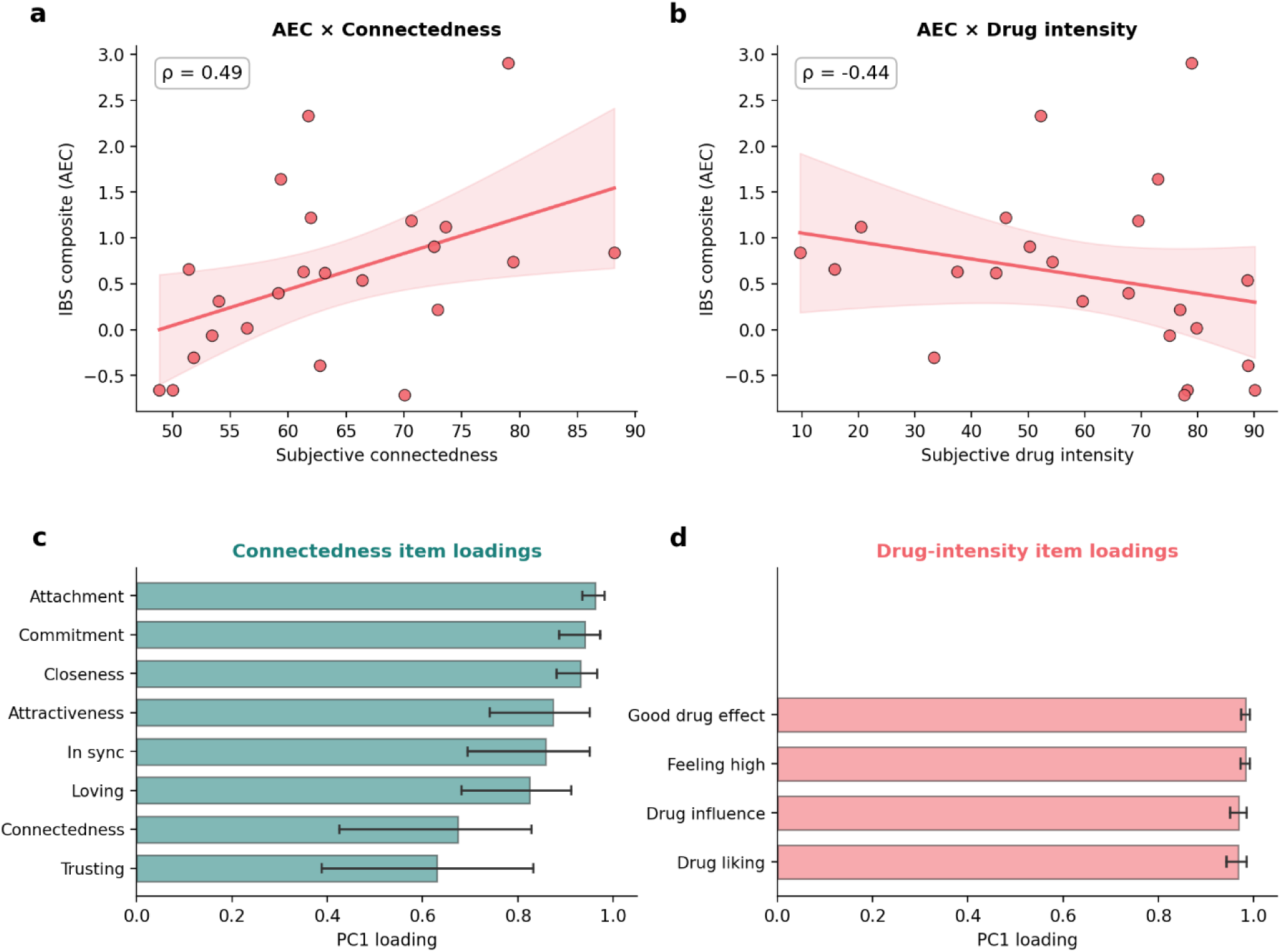
The relationship between resting theta-band amplitude coupling and subjective experience under LSD, and composite item loadings. (a, b) Scatterplots depict the association between the inter-brain synchrony composite (amplitude envelope correlation, AEC) and each visual-analogue-scale composite in the LSD condition: subjective connectedness (a) and subjective drug intensity (b). Each point represents one dyad; the solid line indicates the ordinary-least-squares fit and the shaded band its 95% confidence interval. Associations were quantified using Spearman’s rank correlation (ρ, shown in each panel). (c, d) Loadings of each item on the first principal component of the corresponding composite, shown beneath the scatterplot for the composite it forms: the eight relational items of the Connectedness composite (c, teal) and the four drug-experience items of the Drug-intensity composite (d, coral). Bars show the first-component loading; error bars denote 95% confidence intervals from a cluster bootstrap (5,000 resamples of couples). Full loadings are reported in Tables S2 and S3.

## Discussion

In romantic couples who received LSD together, we found enhanced genuine, partner-specific inter-brain synchrony during rest, broadly increased subjective feelings of connectedness, and, in an exploratory analysis, greater resting synchrony during LSD was linked to stronger perceived connectedness, an effect not observed under placebo. The synchrony effect was specific in three respects: it was carried by theta-band amplitude coupling, it survived two complementary surrogate controls confirming genuine, partner-specific, contemporaneous coupling, and it emerged during rest rather than during structured interaction. To our knowledge, this is the first demonstration that a psychedelic enhances brain-to-brain coupling between interacting people.

The increase in resting theta-band amplitude coupling is consistent with a growing literature implicating theta oscillations in social cognition and interpersonal coordination, including monitoring others, regulating social-emotional responses, and synchronising with partners during interaction(21,31,32). LSD selectively enhanced theta coupling in a partner-specific manner beyond the surrogate baseline, suggesting that the drug amplifies the neural dynamics that ordinarily support attunement between partners rather than producing a non-specific increase in signal similarity. This effect was accompanied by a reorganisation of beta-band phase synchrony, which under LSD lost its normal dependence on eye state. Thus, LSD did not uniformly strengthen all forms of inter-brain coupling, but increased amplitude-based coupling while altering the state-dependence of phase-based coupling, a dissociation consistent with these metrics indexing distinct neural phenomena(26–28).

Why did this coupling emerge at rest rather than during the interactive tasks? One possibility is that LSD enhances inter-brain coupling specifically during unconstrained co-presence, in the absence of any directed task or focus, whereas the structured tasks drew attention toward explicit external demands. The acute effects of psychedelics have previously been found to emerge more readily during spontaneous than goal-directed cognition(33). A similar principle may apply here, whereby partner-specific coupling enhanced by LSD is most readily expressed when cognition is unconstrained by external task demands. More specifically, task performance may divert attention from the subtle interpersonal cues that support mutual attunement. Partners spontaneously adjust their own responses to the affective cues of the other, but when focused on a task, sensitivity to these cues may be reduced (34). Consistent with this, the drawing task constrained coordination around an explicit performance goal, while the eye-contact conditions were behaviourally animated under LSD (for example, partners frequently laughed, often at different moments rather than together) in ways less conducive to sustained mutual attentional engagement. An alternative, not mutually exclusive, account is that the resting recording captured a carry-over of synchrony established during the preceding shared experience under the drug. Consistent with this, inter-brain coupling has been shown to remain elevated during non-interactive periods following a social interaction, and to predict subsequent motivation to connect with the partner(13,35,36). Such post-interaction coupling has been interpreted as a form of inter-brain plasticity, whereby recent shared experience leaves a trace in the coupling between partners’ brains(13). Our design cannot fully determine between these accounts, as both predict partner-specific resting synchrony that correlates with felt connectedness around the resting period. However, the synchrony–connectedness association was strongest at the timepoint immediately adjacent to the resting recording and attenuated thereafter, suggesting that this coupling was meaningfully tied to the resting state rather than reflecting a stable characteristic of the dyad.

Alongside these neural effects, LSD robustly increased subjective interpersonal connectedness, elevating feelings of love, attachment, closeness, trust, and being “in sync,” while reducing loneliness. Related naturalistic work has identified communitas, an acute sense of togetherness and shared humanity, as a potentially important feature of collective psychedelic experiences, associated with subsequent social connectedness and wellbeing(37). Experimental studies have established that psychedelics alter social cognition(7) but this evidence has come almost exclusively from paradigms in which a single participant responds to (non-interactive) stimuli, such as faces or computer-based tasks. Whether and how psychedelics affect social processes between two people actually interacting with one another has remained essentially unexamined. By recording from both partners of a couple as they shared the drug experience, our findings extend this literature from the individual responding in isolation to the dyad in genuine interaction. Critically, this affiliative shift dissociated from the pharmacological state itself: whereas subjective drug intensity and plasma LSD concentrations peaked early and declined over the session, connectedness rose and remained elevated throughout. This temporal divergence indicates that the social effects of LSD are not merely a by-product of feeling intoxicated, and raises the possibility that LSD creates a state in which shared experiences between partners accumulate or exert greater influence on relational feelings over time. However, our design cannot determine which aspects of the shared experience contributed to this trajectory. Converging with this dissociation, greater resting amplitude coupling was associated with lower, rather than higher, subjective drug intensity, the opposite of what a purely pharmacological account of synchrony would predict. The exploratory association between resting amplitude synchrony and connectedness was specific to the LSD condition, absent under placebo, and present for amplitude but not phase coupling, suggesting a degree of specificity in how subjective connection maps onto inter-brain dynamics. These associations did not survive correction for multiple comparisons and should be regarded as hypothesis-generating; nonetheless, its specificity to the drug state and to amplitude coupling makes it a promising target for confirmation in adequately powered designs.

A plausible mechanism begins with 5-HT2A receptor activation. Human blockade studies show that ketanserin attenuates LSD-induced changes in brain connectivity and social processing, implicating this receptor in both its neural and interpersonal effects(38,39). In rodents, LSD increased social behavior through 5-HT2A- and AMPA-dependent excitation of medial prefrontal neurons and downstream mTORC1 signalling, with receptor blockade abolishing these effects(40). LSD also reopened a critical period for social-reward learning through a 5-HT2A-dependent mechanism, and restored impaired inter-brain coupling in dog-human dyads(41,42). Together, these findings suggest that psychedelics like LSD may increase the responsiveness and plasticity of social neural systems, allowing interpersonal experience to more readily shape partner-specific neural dynamics.

Taken together, these findings position inter-brain synchrony as a candidate neural correlate of the social effects of LSD, though not as a straightforward correlate of felt connection. Namely, whereas subjective connectedness was broadly elevated and sustained across the session, partner-specific synchrony emerged only during quiet copresence. This discrepancy may mean that interbrain synchrony reflects neural connection in a specific situation, rather than tracking how connected people feel moment to moment. Indeed, prior work in romantic couples has shown that greater felt support during empathy-giving can coexist with lower inter-brain synchrony, suggesting that established partners may rely more heavily on existing internal models of one another, rather than on moment-to-moment interpersonal coupling(20). That said, the observation that a psychedelic could concurrently heighten the subjective experience of connectedness and enhance a partner-specific, interbrain signature of connection, is noteworthy and offers a potential entry point into the mechanisms by which these drugs reshape social experience. Whereas psychedelics are thought to open transient windows of heightened plasticity within the individual brain(41,43,44), they may also act between brains, transiently increasing the degree to which interacting partners attune to one another, an extension of the notion that interbrain-coupling is itself plastic and shaped by recent social experience(13). One intriguing possibility is that such a state could temporarily loosen established patterns of interpersonal responding, creating greater opportunity for partners to update how they perceive and respond to one another. If so, this combination of heightened connectedness and interpersonal plasticity could have relevance for couple-based therapeutic settings, although this remains to be tested.

Several limitations should be noted: our sample consisted of healthy, established couples, and whether these effects extend to unfamiliar pairs or clinical populations remains to be determined. The association between resting synchrony and connectedness was exploratory, based on a modest sample, and did not survive correction for multiple comparisons; larger studies are needed to confirm this association, though we note that recruiting this population proved highly demanding (Figure S1). A further limitation concerns the integrity of the blind: the subjective effects of LSD meant that participants were frequently able to correctly identify the treatment condition (see supplementary results). As such, functional unblinding cannot be excluded as a contributor to the subjective effects reported here, although it is unclear how this would influence a more objective measure such as inter-brain synchrony. Our EEG approach also captures only one level of interpersonal coupling. Restricting analyses to theta, alpha and beta oscillatory synchrony provided an artefact-resistant measure during naturalistic interaction, but necessarily leaves other neural dynamics unresolved. Future studies combining hyperscanning with richer behavioral and physiological measures, and methods capable of resolving the direction and temporal structure of interpersonal influence, could help determine how neural coupling relates to the moment-to-moment exchange of social information. A final consideration is the timing of neural recording, which was confined to a single window close to the pharmacokinetic peak. This is particularly pertinent given our observation that subjective connectedness remained elevated even as drug intensity declined: synchrony measured after the acute peak, or during the subsequent afterglow in the day(s) following, might yield a different pattern, particularly for interactive states in which acute intoxication could potentially obscure partner-specific coupling. Should inter-brain synchrony prove to be a driver of felt connection rather than merely a correlate, establishing when partner-specific coupling is greatest could inform dyadic therapeutic applications, in which the connection between two people is itself central to the intervention(45,46). Notwithstanding these limitations, by recording simultaneously from two brains sharing a psychedelic state, the present study provides initial evidence that LSD enhances genuine inter-brain synchrony between people, linking a pharmacologically induced state of felt connection to a measurable signature shared across interacting brains.

## Acknowledgements

This study was supported by the Mind Science Foundation (2023) Brainstorm award (to NLM, PM, JGR). NLM is financially supported by the Dutch Research Council (NWO, grant number VI.Veni.231G.011). GD is supported by the Institute for Data Valorization, Montreal and the Canada First Research Excellence Fund (IVADO; CF00137433), the Fonds de recherche du Québec (FRQ; 285289), the Natural Sciences and Engineering Research Council of Canada (NSERC; DGECR-2023-00089). During the preparation of this work, the author(s) used Claude (Anthropic) to refine, edit, and format human-written text; to produce code used to generate figures and analyze data; and to format tables. All AI-assisted output was reviewed, verified, and edited by the author(s), who take full responsibility for the content of the publication. We are deeply grateful to all participants for their time, effort, and trust, and to Cees van Leeuwen for providing medical supervision. We also thank the interns who contributed to participant recruitment, screening, and data collection, including Nina Stoel, Margherita Rigoni, Jonas Neubert, Logan Riffey, Brix Binek, Anna Gaidosch, Igor Esteban Gamez, Alissa Haj Yahya, Milena Heibrock, Ana-Alexandra Luta, Lea McQuaid, Szabolcs Meszaros, Bent Mucke, Julia Nowicka, Jack Pahlevan, Max Payton, Max Reim, Jenelle Rofe, and Petros Tsiakas. We thank Bram Terstappen for his invaluable logistical support throughout the data collection period, and Johan Gielissen for setting up and troubleshooting the hyperscanning system.

## Data availability

Data can be made available upon reasonable request to N.L.M. and a data use agreement executed with Maastricht University.

## Supplementary Information for

**Figure S1.**
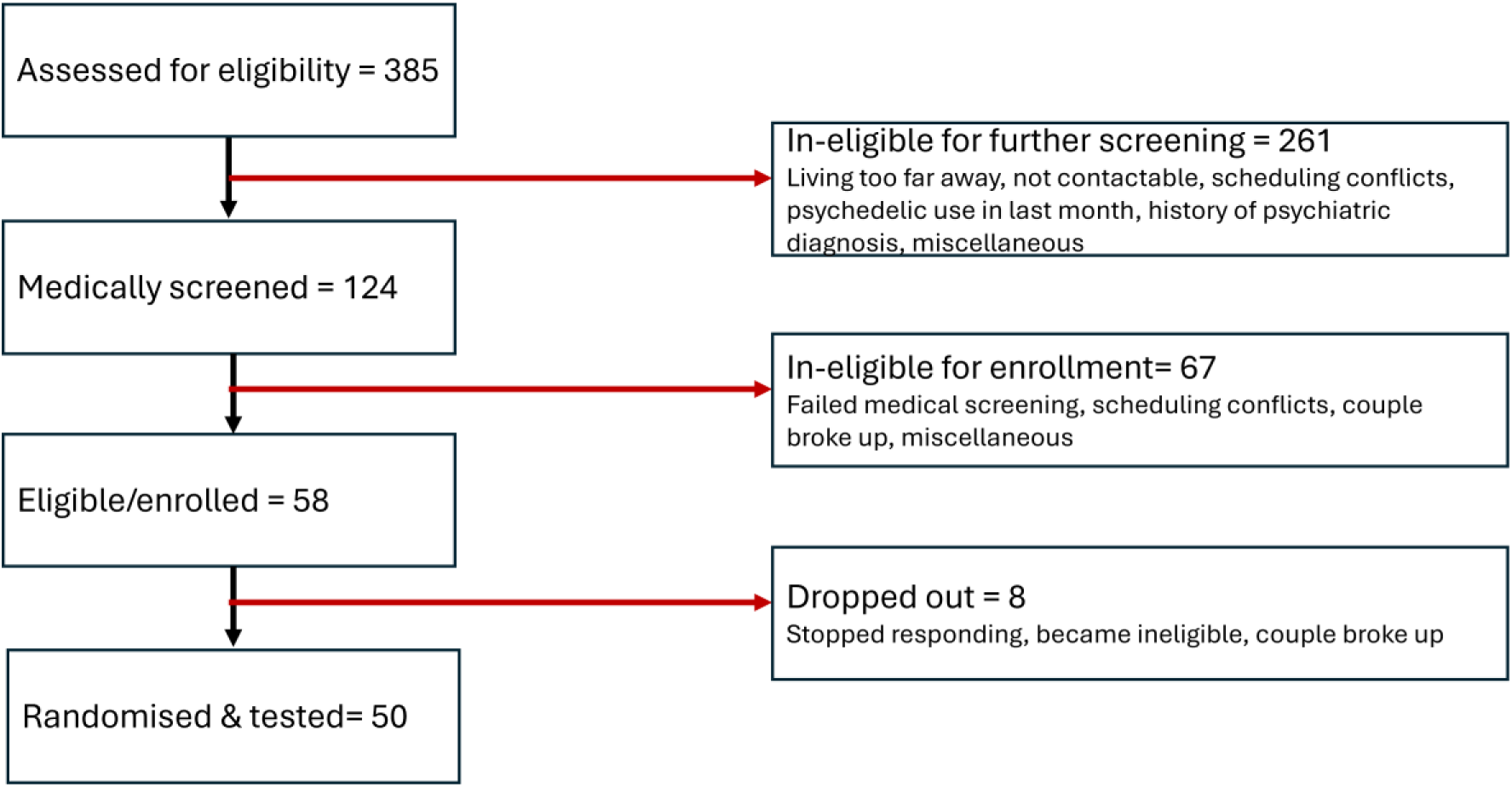
Recruitment overflow.

**Figure S2.**
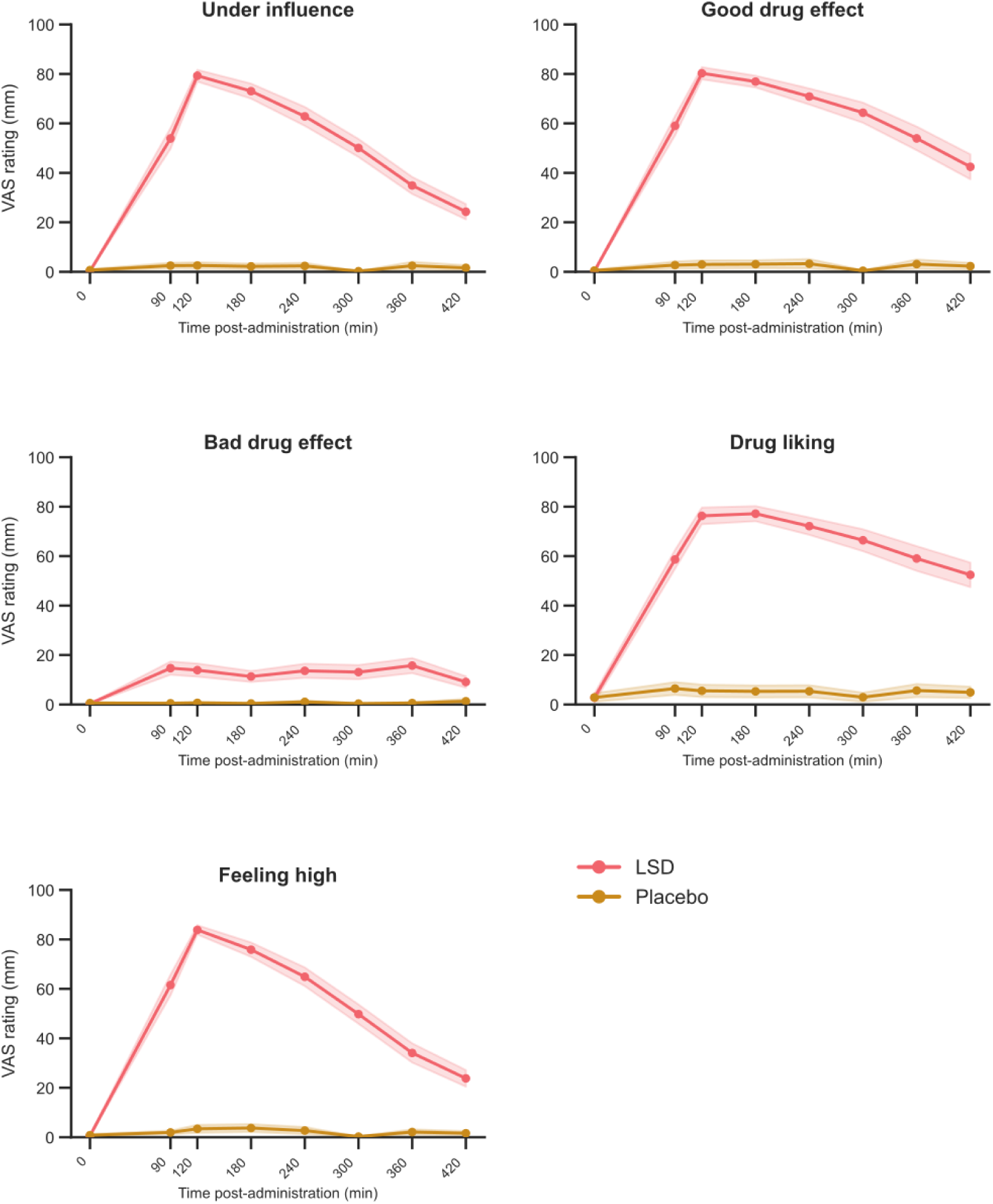
Subjective drug-experience ratings across the testing day under LSD and placebo. Mean visual analogue scale (VAS) ratings for the drug-experience items at eight assessment timepoints (0–420 min post-administration) under LSD (coral) and placebo (gold). Ratings are on a 0–100 mm scale. Shaded bands denote ±1 standard error of the mean. Data are shown at the individual-participant level (N = 50)

**Figure S3.**
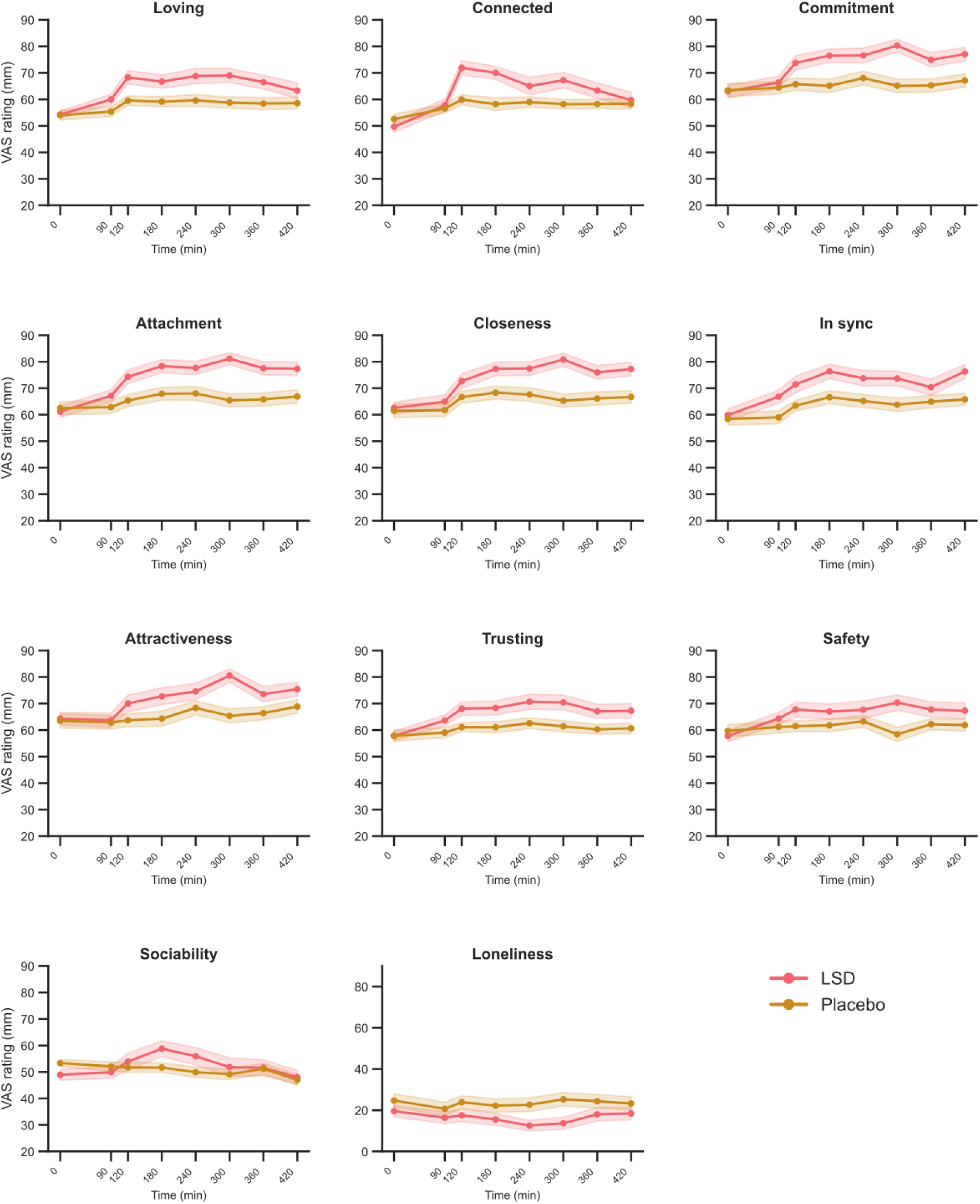
Subjective relational ratings across the testing day under LSD and placebo. Mean visual analogue scale (VAS) ratings for eleven relational items at eight assessment timepoints (0–420 min post-administration) under LSD (coral) and placebo (gold). Shaded bands denote ±1 standard error of the mean. For visualisation, items are displayed on a 20–90 mm range (loneliness, 0–90 mm) to highlight condition differences; all ratings were made on a 0–100 mm scale, with the midpoint (50) anchored to participants’ “normal” level for each construct. Data are shown at the individual-participant level (N = 50).

**Figure S4.**
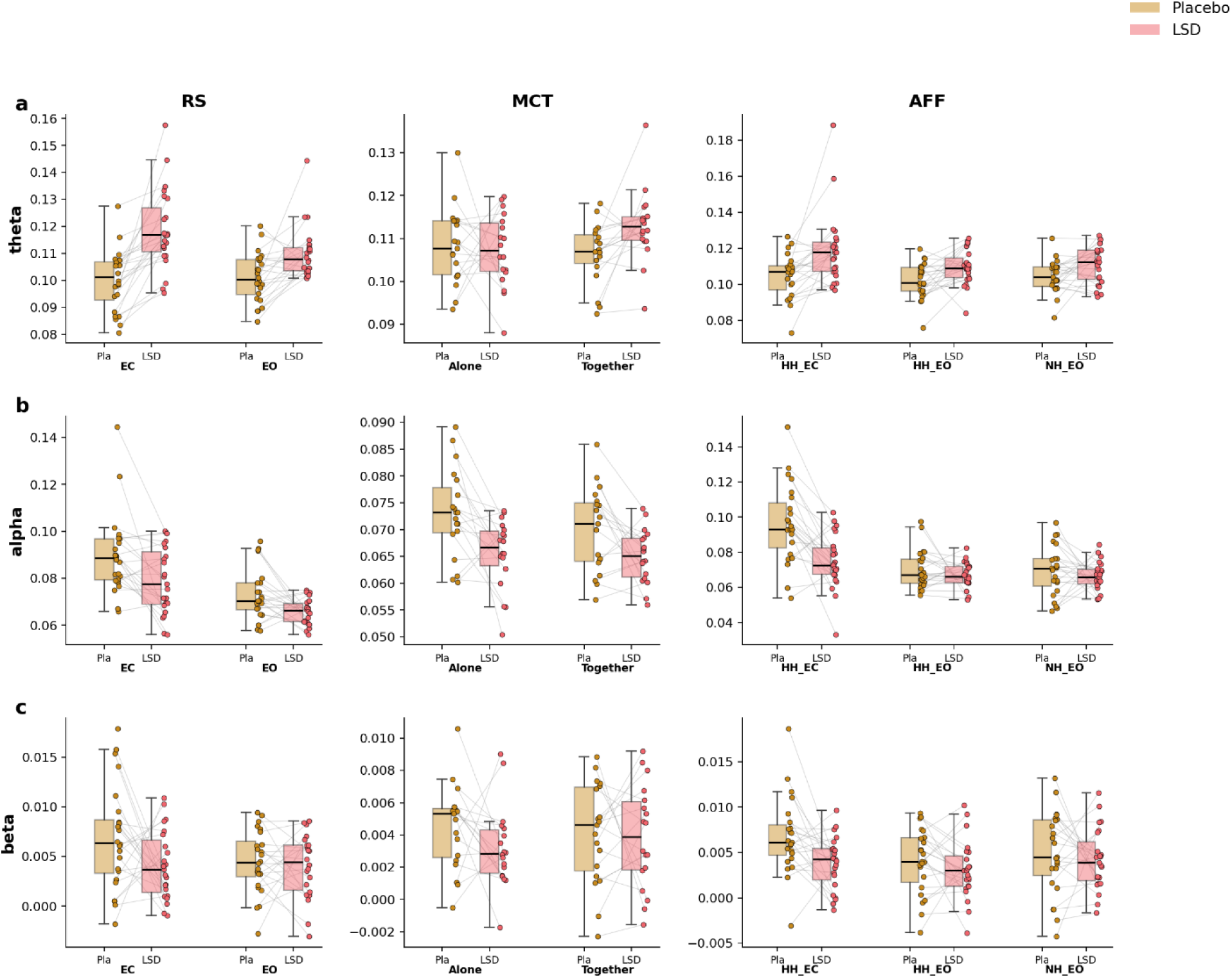
Task-averaged inter-brain amplitude coupling (AEC) across frequency bands and tasks. Inter-brain synchrony was computed for each dyad as the mean AEC across all fifteen region-of-interest (ROI) pairings among the five electrode clusters (frontal, fronto-central, central, parieto-temporal, occipital), yielding a single value per couple, condition, and drug session. Rows correspond to frequency band, (a) theta, (b) alpha, (c) beta, and columns to task (resting state, RS; motor cooperation, MCT; affective touch, AFF). Within each panel, data are split by drug condition (placebo, gold; LSD, coral) and by the conditions specific to each task (RS: eyes closed, EC / eyes open, EO; MCT: Alone / Together; AFF: hand-holding eyes-closed, HH_EC / hand-holding eyes-open, HH_EO / no-touch eyes-open, NH_EO). Boxplots depict the median and interquartile range with whiskers extending to the furthest data points within 1.5×IQR of the quartiles; overlaid points represent individual dyads, and thin grey lines connect each dyad across the placebo and LSD sessions.

**Figure S5.**
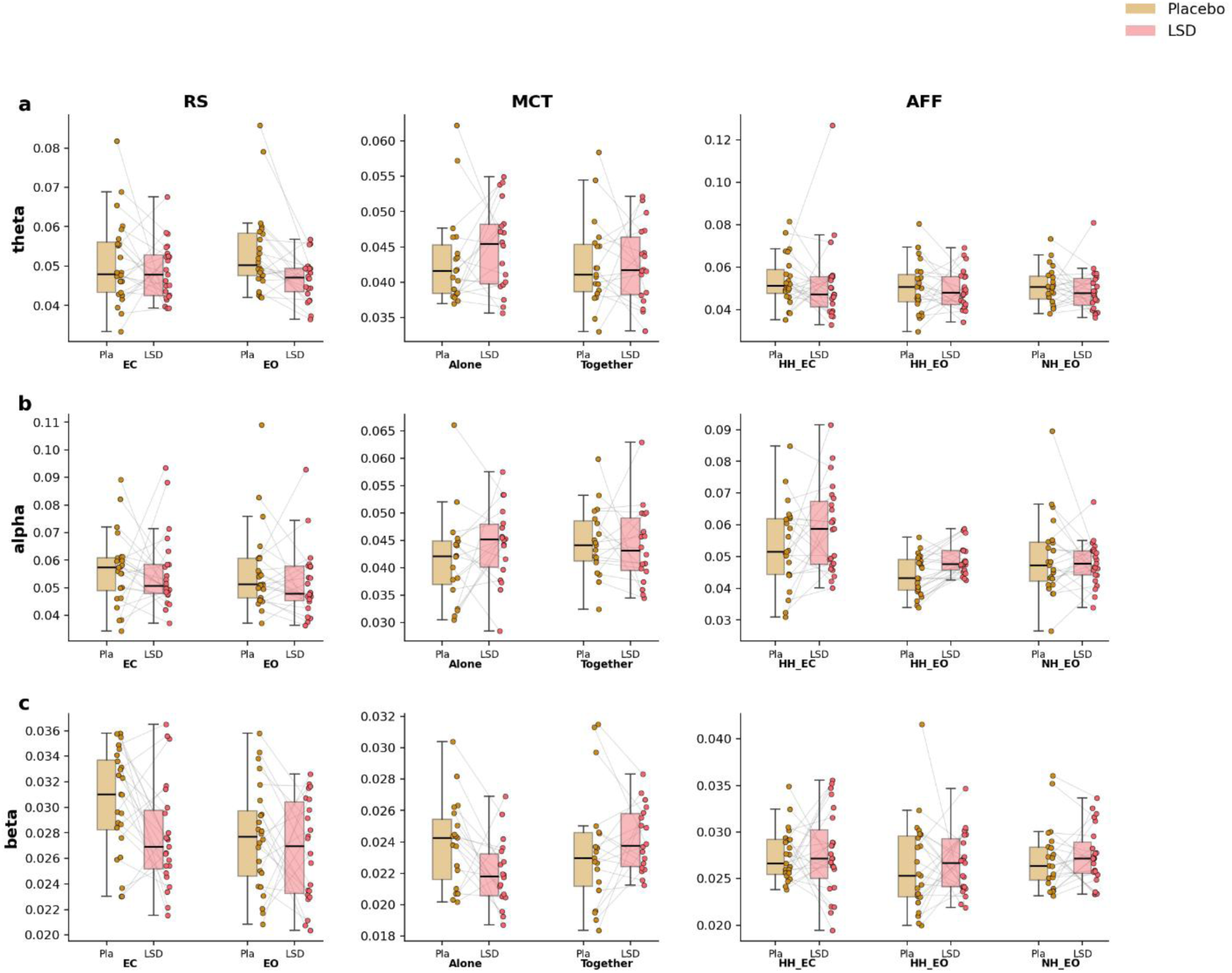
Task-averaged inter-brain phase synchrony (wPLI) across frequency bands and tasks. Inter-brain synchrony was computed for each dyad as the mean wPLI across all fifteen region-of-interest (ROI) pairings among the five electrode clusters (frontal, fronto-central, central, parieto-temporal, occipital), yielding a single value per couple, condition, and drug session. Rows correspond to frequency band, (a) theta, (b) alpha, (c) beta, and columns to task (resting state, RS; motor cooperation, MCT; affective touch, AFF). Within each panel, data are split by drug condition (placebo, gold; LSD, coral) and by the conditions specific to each task (RS: eyes closed, EC / eyes open, EO; MCT: Alone / Together; AFF: hand-holding eyes-closed, HH_EC / hand-holding eyes-open, HH_EO / no-touch eyes-open, NH_EO). Boxplots depict the median and interquartile range with whiskers extending to the furthest data points within 1.5×IQR of the quartiles; overlaid points represent individual dyads, and thin grey lines connect each dyad across the placebo and LSD sessions.

**Table S1.** Participant demographics. Values are mean (SD) for continuous variables and n (%) for categorical variables. Alcohol and caffeine refer to current typical consumption; MDMA and psychedelic use refer to lifetime occasions.

| Characteristic | Value |
| --- | --- |
| **Demographics** |  |
| Age (years) | 25.5 (5.2) |
| Gender, women | 27 (54.0%) |
| Gender, men | 21 (42.0%) |
| Gender, non-binary | 2 (4.0%) |
| **Couple composition** |  |
| Mixed-gender | 21 (84.0%) |
| Female same-gender | 3 (12.0%) |
| Non-binary | 1 (4.0%) |
| **Highest education** |  |
| Secondary or vocational | 19 (38.0%) |
| Bachelor | 20 (40.0%) |
| Master's or higher | 11 (22.0%) |
| **Substance use** |  |
| Alcohol (units/week) | 2.4 (3.1) |
| Caffeine (units/day) | 1.2 (1.0) |
| MDMA (lifetime occasions) | 5.0 (8.0) |
| Psychedelics (lifetime occasions) | 9.5 (12.6) |
| **Relationship length (per couple)** |  |
| 6 months to 1 year | 4 (16.0%) |
| 1 to 2 years | 10 (40.0%) |
| More than 2 years | 11 (44.0%) |

**Table S2.** Factor loadings of the Connectedness composite. Loadings on the principal component analysis of the eleven relational visual analogue scale items. Eight items loaded strongly on the first component (.67–.95), clearly separated from the remaining items, which loaded ≤ |.51|; these eight items formed the Connectedness composite (Cronbach’s α = .94).

| Item | Component |  |  |
| --- | --- | --- | --- |
|  | 1 | 2 | 3 |
| Retained in the Connectedness composite |  |  |  |
| Attachment | .950 |  |  |
| Commitment | .932 |  |  |
| Closeness | .922 |  |  |
| In sync | .853 |  |  |
| Attractiveness | .853 |  |  |
| Loving | .822 |  |  |
| Connectedness | .679 |  |  |
| Trusting | .666 | .670 |  |
| Excluded |  |  |  |
| Safety | .500 | .766 |  |
| Loneliness | -.513 |  |  |
| Sociability |  |  | .858 |

**Table S3.** Factor loadings of the Drug-intensity composite. Loadings on the principal component analysis of the five drug-experience visual analogue scale items. Four items loaded strongly on the first component (.95–.98), clearly separated from Bad drug effect, which loaded ≤ |.62| on the first component and split onto a distinct second component; these four items formed the Drug-intensity composite (Cronbach’s α = .96).

| Item | Component |  |
| --- | --- | --- |
|  | 1 | 2 |
| Retained in the Drug-intensity composite |  |  |
| Feeling high | .983 |  |
| Drug influence | .973 |  |
| Good drug effect | .965 |  |
| Drug liking | .948 |  |
| Excluded |  |  |
| Bad drug effect | .621 | .781 |

**Table S4.** Effect sizes for the condition difference on each VAS item. *Effect* names whichever term is significant in the model; where the Treatment × Time interaction is significant the time-varying effect is the one listed, otherwise the Treatment effect is. η²*p* was derived from the *F* statistic and its degrees of freedom, and refers to the term named in the *Effect* column. *Δ* and *d* are the estimated marginal mean difference between conditions (LSD − placebo) averaged across the eight assessment times, in VAS points (0–100) and in standard deviation units respectively; *d* was obtained by dividing Δ by the model-implied total between-person standard deviation, the square root of the sum of the variance components. The confidence interval is for *d*, obtained by expressing the confidence limits for Δ on the same scale. *Peak* gives the largest single-timepoint *d* with its time in minutes, shown only where the interaction is significant. Positive values indicate higher ratings under LSD. Note that η²ₚ refers to the term named in the Effect column and is therefore not comparable across rows, whereas Δ and *d* are.

| Item | Effect | $\eta^2_p$ | $\Delta$ | *d* | 95% CI | Peak |
| --- | --- | --- | --- | --- | --- | --- |
| **Relational items** |  |  |  |  |  |  |
| Connectedness | Treatment × Time | 0.05 | 5.8 | 0.36 | [0.19, 0.52] | 0.76 (120) |
| Attachment | Treatment × Time | 0.05 | 9.0 | 0.54 | [0.37, 0.70] | 0.86 (300) |
| Commitment | Treatment × Time | 0.05 | 8.4 | 0.48 | [0.31, 0.65] | 0.81 (300) |
| Safety | Treatment × Time | 0.04 | 5.5 | 0.32 | [0.16, 0.47] | 0.65 (300) |
| Closeness | Treatment × Time | 0.03 | 8.6 | 0.50 | [0.36, 0.64] | 0.83 (300) |
| Attractiveness | Treatment × Time | 0.03 | 6.6 | 0.35 | [0.21, 0.50] | 0.69 (300) |
| Loving | Treatment | 0.22 | 7.1 | 0.46 | [0.30, 0.62] | — |
| In sync | Treatment | 0.21 | 8.0 | 0.46 | [0.31, 0.60] | — |
| Trusting | Treatment | 0.21 | 6.7 | 0.43 | [0.28, 0.58] | — |
| Loneliness | Treatment | 0.15 | -7.3 | -0.34 | [-0.49, -0.19] | — |
| Sociability | — | — | 1.5 | 0.09 | [-0.09, 0.27] | — |
| **Drug-intensity items** |  |  |  |  |  |  |
| Under the influence | Treatment × Time | 0.50 | 45.4 | 2.73 | [2.53, 2.94] | 4.63 (120) |
| Feeling high | Treatment × Time | 0.53 | 47.1 | 2.79 | [2.58, 3.00] | 4.77 (120) |
| Good drug effect | Treatment × Time | 0.47 | 53.7 | 2.78 | [2.54, 3.01] | 4.01 (120) |
| Drug liking | Treatment × Time | 0.40 | 53.3 | 2.39 | [2.14, 2.64] | 3.25 (180) |
| Bad drug effect | Treatment × Time | 0.09 | 10.5 | 0.85 | [0.64, 1.06] | 1.20 (360) |

## Supplement

### Participants

Volunteers had to fulfill a number of general inclusion and exclusion criteria before being admitted to the study, with both partners having to fulfill all of the criteria. Main inclusion criteria included previous experience with a psychedelic, absence of any major medical or psychological conditions as determined by a medical examination and laboratory analysis, no psychedelic use in the past 3 months, and willingness to refrain from taking illicit psychoactive substances during the study.

Healthy participants were recruited, who met the following criteria: 18–40 years old; previous experience with a psychedelic drug, but not within the past 3 months; normal weight, body mass index between 18 and 28 kg/m2; free from psychotropic medication; good physical health, including absence of major medical, endocrine, and neurological conditions; willingness to refrain from taking illicit psychoactive substances during the study; and written informed consent. Exclusion criteria consisted of: history of drug abuse or addiction; pregnancy or lactation; health issues including hypertension (diastolic > 90 and systolic > 140), cardiac dysfunction, and liver dysfunction; current psychiatric disorder; previous experience of serious side effects to psychedelics. Before inclusion, subjects answered medical questionnaires about their health and drug use, and were screened and examined by a study physician, who checked for general health, conducted a resting ECG, and took blood and urine samples in which hematology, clinical chemistry, urine, and virology analyses were conducted. Participants were financially compensated for their participation in the study.

### Drug

LSD (free-base solution) doses were prepared by dissolving the compound in ethanol (96% vol). The placebo solution consisted of ethanol alone (1 ml; Holze et al., 2020). On each testing day, participants received 2 ml of liquid solution (LSD in ethanol or ethanol only) administered orally.

### Randomization and blinding

An experimenter who was not responsible for treatment randomization or preparation, recruited all participants. A separate experimenter, who did not come in direct contact with the subjects, allocated treatment conditions. LSD and placebo were administered in an AB or BA sequence, with couples randomly allocated to treatment order such that half followed each sequence. Randomization was performed using Sealed Envelope software by an experimenter who had no direct contact with participants. The study was double-blind, with both participants and testing experimenters blinded to treatment conditions. LSD was prepared by a separate experimenter who was not involved in participant testing.

### Sample Size

Power analysis for neuroimaging data is complex because analyses involve many non-independent comparisons, and statistical power varies across brain regions(47). Accordingly, sample sizes are often informed by previous empirical work in the field. Previous hyperscanning studies using comparable paradigms have detected inter-brain neural synchrony in samples of 18–23 couples(17,20), while neuroimaging studies of psychedelic drug effects have commonly employed samples of 10–20 participants(48–50). Based on this prior work, we expected that a sample of N = 25 couples (50 participants) would provide sufficient power to detect drug-related effects on inter-brain synchrony.

### Procedures

A training session took place before the test days in order to familiarize participants with the tests and test procedures. Participants had to refrain from psychedelic substance use three months before study start, and from other drug use at least one week before study start, until completion of all testing days. Participants were requested to not consume caffeinated or alcoholic beverages after midnight of the evening before the test days, as well as during the test days. On the test days, smokers were asked to refrain from nicotine use 2 h before the start of the test days as well as during the testdays. All participants were expected to arrive well rested at the test facilities at 9 AM.

Participants spent the entire day in a room, set up like a living room setting, equipped with a couch, table, chairs, and lamps. All measurements were conducted in this room. At arrival, participants were screened for the presence of alcohol in breath, drugs of abuse in urine (THC/ opiates/ cocaine/ amphetamines/ methamphetamines), and women were tested for pregnancy. When tests were negative, a cannula was placed in order to take blood throughout the testing day, and subjective measures were taken (visual analogue scales). At 10 AM, drug administration took place, and participants were fitted with their own EEG cap, electrodes, and ECG electrodes to measure heart rate. One hour to 8 hours post-administration, participants completed a test battery consisting of questionnaires, EEG measures, and blood samples (Figure 1).

## Methods

### LSD concentrations

Blood samples for plasma concentration were centrifuged and plasma was frozen at −30°C until analysis. Analysis of LSD was via serum (500 µl), extracted with 2.5 ml of 1-chlorobutane/diethyl ether (50:50, v/v) after addition of 0.5 mL 1M phosphate buffer pH 9.5 and 50 µl of internal standard (acetonitrile containing 2.5 ng of LSD-d3 from Cerilliant, Round Rock, Texas, US). The organic phase was evaporated and reconstituted with 100 µl of 0.1 % formic acid/acetonitrile (80:20, v/v). The analysis of 5 µl was performed on an Agilent (Waldbronn, Germany) LC-MS/MS system consisting of a 1290 Infinity II Liquid Chromatograph coupled via JetStream Electrospray Interface (ESI) to a G6495D Triple Quadrupole Mass Spectrometer. Analytes were separated on a Kinetex® 2.6 µm XB-C18 100 Å LC column (100 x 2.1 mm) plus corresponding guard column from Phenomenex (Aschaffenburg, Germany) at 50 °C. Gradient elution at a flow rate of 0.5 ml/min using 0.01% formic acid containing 5 mM ammonium formate (A) and acetonitrile containing 0.1 % formic acid (B) started with 20 % B, increased to 50 % B during 1.5 min and further to 100 % B during 2.5 min. Source parameters were: gas temperature 250 °C, gas flow 13 l/min, nebulizer 20 psi, sheath gas temperature 400 °C, sheath gas flow 12 l/min and capillary voltage 4000 V. Detection was performed in the multiple reaction monitoring mode (m/z, collision energy in parentheses, quantifier underlined): LSD-d3: 327.2→226.1 (28V); LSD 324.2→208.1 (32V); 223.1 (28V). Six calibration standards were prepared in the range 5 – 2500 pg per ml human serum and were analysed with the samples. Calibrations were linear (regression coefficients >0.99).

## Statistics

### Composite construction and reliability

Reliability of all composites was assessed with Cronbach’s α computed on the couple-level scores. The exploratory PCA of the relational VAS items (Kaiser-Meyer-Olkin = .84; Bartlett’s test of sphericity p < .001) produced a dominant first component accounting for 56% of variance, on which eight items loaded strongly (.67–.95): loving, connected, commitment, attachment, close, in-sync, attractive, and trusting. These were averaged into a Connectedness composite (α = .94). The remaining relational items (sociability, loneliness, safety) did not load on this component and were not included in the composite.

A PCA of the five drug-experience items yielded a dominant first component (83% of variance) on which four loaded strongly (.95–.98): drug influence, good drug effect, drug liking, and feeling high. These formed the Drug-intensity composite (α = .96). Bad drug effect was excluded, loading weakly and splitting onto a separate component (Table S3).

On the neural side, only the connectivity metrics and ROI pairs showing a significant effect of Treatment were considered; all four amplitude envelope correlation (AEC) ROI pairs and both weighted phase-lag index (wPLI) ROI pairs met this criterion. Within each metric the ROI-pair measures were highly intercorrelated (AEC: r = .83–.96, α = .98; wPLI: α = .83) and were z-standardized across the full sample before averaging into an AEC composite and a wPLI composite. The two metrics were analyzed separately, as phase and amplitude coupling index distinct connectivity phenomena; consistent with this, the AEC and wPLI composites were uncorrelated (r = −.09, p = .55).

### Partner concordance and couple-level aggregation

Because interbrain synchrony is a dyadic measure, partner VAS ratings were averaged into couple-level scores. To evaluate this aggregation, we examined partner concordance at the analyzed timepoint using two-way random, single-measures intraclass correlations. Concordance was strong for drug intensity (ICC = .90) and moderate but statistically reliable for connectedness (ICC = .42, p = .001), supporting couple-level aggregation for both measures.

## Results

### Pharmacokinetic timecourse

The maximum mean LSD serum concentration (Cmax) was 668.96 pg/mL (68.20-1439.26, n=35), concordant with the applied oral dose of 50 μg. The mean elimination half-life was 3.64 hours (1.70-6.09, n=24), in line with previous work(51–53). Figure 2 shows the overall pharmacokinetic timecourse.

### Temporal-shift surrogate control for resting-state theta-AEC

To assess whether the resting-state theta-AEC effects could be explained by shared pharmacodynamic or vigilance nonstationarity, rather than genuine moment-to-moment inter-brain coupling, we applied an additional within-couple temporal surrogate. For each RS session, one partner’s epoch order was circularly shifted by a random integer amount within the eyes-open or eyes-closed block, and AEC was recomputed across 1,000 shifts. This procedure preserves the pharmacological context (both partners remain within the same ∼8-minute recording block under the same drug condition) while destroying the fine-grained temporal alignment required for genuine synchrony. All four theta-AEC ROI pairs exceeded the temporal-shift surrogate baseline under both LSD and Placebo (empirical p = 0.001 for all 8 drug × ROI-pair combinations; N = 22 dyads), confirming that the observed effects depend on contemporaneous amplitude-envelope alignment and are not an artefact of shared pharmacodynamic nonstationarity.

### No relationship between neural synchrony and VAS items under placebo

To assess whether the LSD findings were state-specific, the primary correlation analysis was repeated within the placebo condition (N = 23). The AEC–Connectedness association observed under LSD was absent under placebo (ρ = −.05, p = .802). Correlations involving subjective drug intensity were not computed for the placebo condition, as drug-intensity ratings were at floor under placebo (M ≈ 3 on a 0–100 scale) and lacked the variance required for meaningful correlation. The wPLI composite was unrelated to Connectedness under placebo (ρ = −.03, p = .890). The absence of the AEC–Connectedness association under placebo indicates that the relationship between interbrain amplitude coupling and subjective connectedness was specific to the LSD state.

### Temporal specificity of the synchrony–connectedness association

To probe the temporal specificity of the resting-state association between inter-brain amplitude coupling (AEC) and subjective connectedness, we examined this association at two timepoints across the session. This exploratory analysis was conducted within the LSD condition at the couple level using Spearman correlations and was not corrected for multiple comparisons. Subjective connectedness was assessed immediately before the resting-state recording (T1) and later in the session, after the two interactive tasks (affective touch and motor cooperation; T2). The association between resting AEC and connectedness was strongest at the resting-adjacent timepoint and was descriptively weaker at the later assessment (T2: ρ = .29, p = .180). The inverse association with subjective drug intensity remained at the later timepoint (ρ = −.44, p = .035). This pattern is consistent with the synchrony–connectedness association being more closely tied to the resting-state period than to a stable, time-invariant characteristic of the dyad.

### Blinding integrity and psychedelic advocacy

To assess the integrity of the blind, participants were asked at the end of each dosing day to guess which treatment they had received, along with a rating of how certain they were (0-100%). Across the 98 possible sessions (48 participants completing both sessions, and 2 participants completing one session), a treatment guess was provided on 95 occasions (n=3 missing). Of the 95 guesses, 93 (97.9%) correctly identified the treatment condition (LSD 100% correct, placebo 95.8% correct). Reported certainty was consistently high (median = 100%; 80 of 95 guesses made with 100% certainty).

Upon study enrollment, participants were asked to describe their attitudes toward psychedelics(54). 76% of participants stated that psychedelics have potential to be beneficial under most circumstances, 18% stated psychedelics have equal potential to be harmful or beneficial, 6% stated that psychedelics are almost always beneficial, 0% stated that psychedelics are almost always harmful or that psychedelics have potential to cause harm under most circumstances.

